# Akt1 Nitration Promotes Proliferation and Mesenchymal Transition Exacerbating Pulmonary Hypertension

**DOI:** 10.1101/2025.10.26.684547

**Authors:** Mathews V Varghese, Dinesh Bharti, Joel James, Paul R. Langlais, Olga Rafikova, Ruslan Rafikov

## Abstract

Pulmonary arterial hypertension (PAH) is a progressive disease characterized by vascular remodeling and increased pulmonary arterial resistance. This study investigates the role of Akt1 nitration in PAH development, focusing on its effect on endothelial-to-mesenchymal transition (EndMT) and vascular cell proliferation. Using the novel Akt1^Y350F^ mutant mouse model, which resists nitration due to a tyrosine-to-phenylalanine substitution, we demonstrated that Akt1 nitration is a key pathogenic factor in PAH progression. Our results show that Akt1^Y350F^ mice, resistant to Akt1 nitration, are protected against PAH in a SU5416/hypoxia (SU5416/Hx) model, exhibiting lower right ventricular systolic pressure (RVSP), reduced right ventricular hypertrophy, and decreased vascular occlusion. Additionally, we identified important molecular mechanisms involving TWIST1, αSMA, HIF1α, and STAT3 signaling pathways that influence EndMT and vascular remodeling. The proteomic analysis revealed other affected pathways, including angiogenesis, lipid metabolism, and mitochondrial function. We demonstrate that oxidative and nitrative stress-induced post-translational modifications contribute to the pathological processes leading to pulmonary hypertension, using a unique Akt nitration-resistant mouse model. These findings provide new insights into the molecular mechanisms underlying PAH and suggest that targeting Akt1 nitration could be a promising therapeutic approach for this devastating disease.

## Introduction

Pulmonary arterial hypertension (PAH) is a progressive and life-threatening disease characterized by elevated pulmonary vascular resistance due to proliferating vascular cells, resulting in right ventricular failure. Despite advances in treatment options, PAH continues to have a poor prognosis, with a 5-year survival rate of approximately 50% ^1^. The pathogenesis of PAH is complex, involving multiple cellular and molecular mechanisms that lead to vascular remodeling, vasoconstriction, and thrombosis ^2^. Different types of endothelial dysfunction play a crucial role in the initiation and progression of PAH ^3^. One of the key processes in vascular remodeling is endothelial proliferation and mesenchymal transition (EndMT), where endothelial cells lose their function and characteristic markers and acquire mesenchymal-like properties ^2^. This phenotypic switch contributes to the abnormal proliferation of vascular cells and the occlusion of pulmonary arterioles, hallmarks of PAH pathology ^4^.

The serine/threonine kinase Akt, particularly its isoform Akt1, is a critical regulator of endothelial cell function and vascular homeostasis ^5^. Under normal conditions, Akt signaling promotes endothelial cell survival, metabolism regulation, proliferation, and nitric oxide production ^6^. However, the role of Akt in PAH pathogenesis remains debated, with some studies implicating the hyperactivation of Akt in disease progression ^7^. Recent evidence has highlighted the importance of post-translational modifications in modulating protein function and cellular signaling. Nitrosative/nitrative stress, characterized by elevated levels of reactive nitrogen species such as peroxynitrite (ONOO-), has been implicated in various cardiovascular diseases, including PAH ^8,9^. Protein tyrosine nitration, a covalent modification induced by peroxynitrite, can significantly alter protein structure and function ^10^. The potential role of Akt1 nitration in PAH pathogenesis has emerged as an interesting area of investigation. Previous studies have shown that Akt can be nitrated at specific tyrosine residues, leading to changes in its activity ^11,12^. However, the particular impact of Akt1 nitration on endothelial function and its contribution to PAH development remains largely unexplored. Understanding the molecular mechanisms of PAH progression is crucial for developing more effective therapeutic strategies. Given the central role of Akt signaling in vascular biology and the increasing recognition of nitrosative stress in PAH, we hypothesized that Akt1 nitration might be a critical driver of endothelial dysfunction and vascular remodeling in PAH.

In this study, we aimed to elucidate the role of Akt1 nitration in PAH pathogenesis, focusing on its impact on EndMT and vascular cell proliferation. We utilized the Akt1^Y350F^ mutant mouse model, which is resistant to nitration at a critical tyrosine residue, to investigate the effects of preventing Akt1 nitration in an established model of PAH. Our findings provide new insights into the molecular pathogenesis of PAH and suggest that targeting Akt1 nitration may represent a novel therapeutic approach for this devastating disease.

## Results

Figure 1 reports the evidence for the role of Akt1 nitration in the pathogenesis of pulmonary arterial hypertension (PAH) in a SU5416/hypoxia (Su/Hx) mouse model and comparing wild-type (WT) mice with Akt1^Y350F^ mutant mice. Western blot analysis of nitrated Akt (Y350NO2-Akt antibody) levels in lung tissue from WT and Akt1^Y350F^ mice shows that the Su/Hx model significantly increased N-Akt levels by 1.5-fold. Akt1^Y350F^ mice showed background nitrated-Akt levels, both with and without Su/Hx treatment (Fig.1A). The stain-free total protein measurement is a loading control, ensuring equal protein loading across samples ^13^. These results demonstrate that the Su/Hx PAH model induces Akt1 nitration and that the Y350F mutation effectively prevents this nitration.

**Figure 1.**
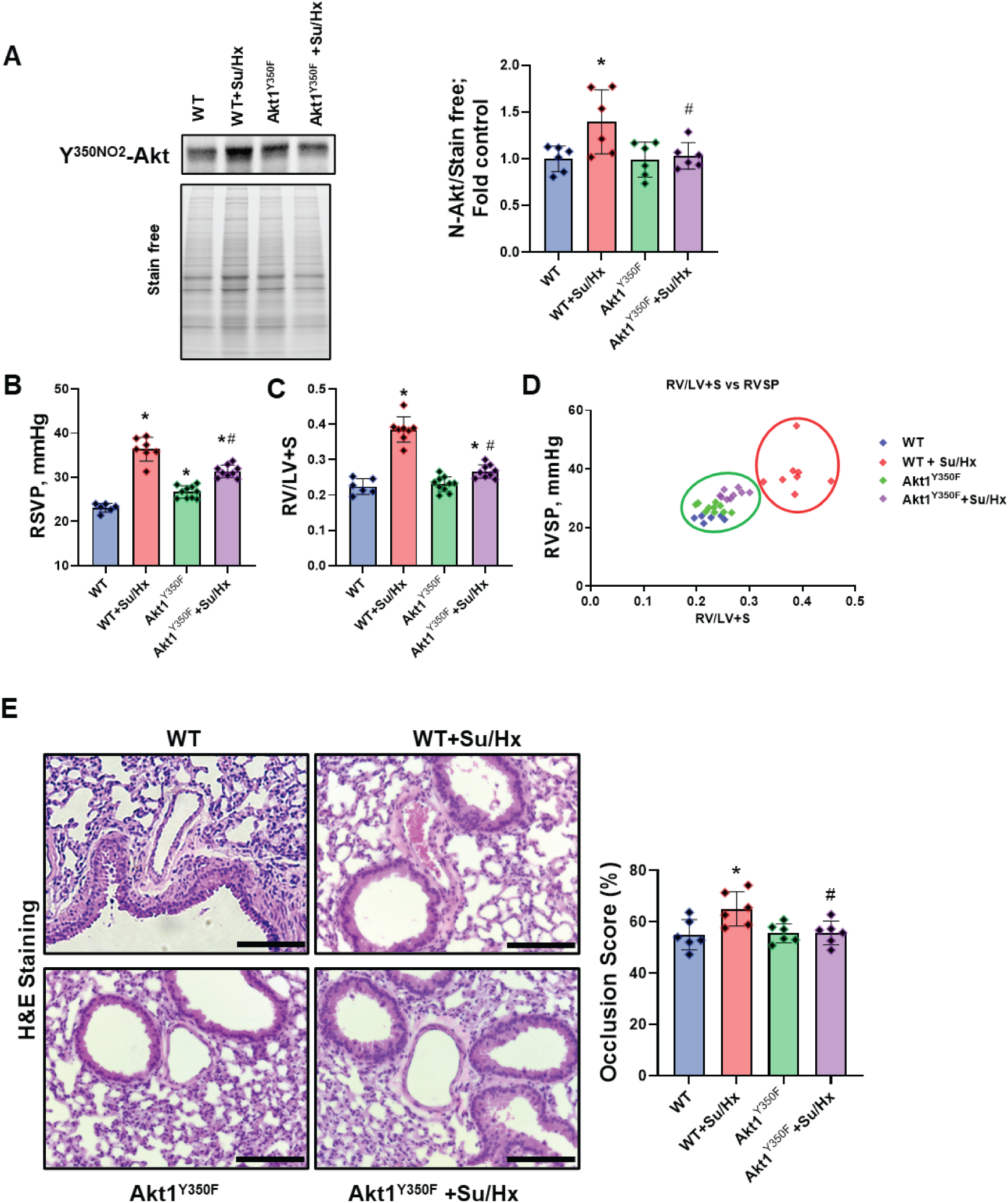
Role of Akt Nitration in PAH: Akt nitration was found to be increased in the wild-type (WT) Sugen hypoxia (Su/Hx) PAH model (Figure 1A). WT mice treated with Su/Hx showed an increase in RVSP (right ventricular systolic pressure) (Figure 1B) and the Fulton index, which is the ratio of the right ventricle to the left ventricle plus septum (Figure 1C). Additionally, correlation analysis between RVSP and the Fulton index showed a significant separation in the WT PAH model compared to the other groups (Figure 1D). In contrast, the Akt1^Y350F^ PAH model significantly attenuated PAH development. Histopathological analysis also showed the Akt1^Y350F^ model provided protection against vascular remodeling caused due to the Su/Hx treatment (Figure 1E). Data are represented as Mean±SD, N=6-9, *p<0.05 vs. WT, ^#^p<0.05 vs. WT Su/HX by 1-way ANOVA with Bonferroni multiple comparison test.

WT mice treated with Su/Hx showed elevated RVSP (Fig.1B) (∼35-40 mmHg compared to 20-25 mmHg in untreated WT, p < 0.05). Akt1^Y350F^ mice treated with Su/Hx maintained significantly lower RVSP than WT mice (p < 0.05). WT mice treated with Su/Hx showed a significant increase in RV/LV+S ratio, whereas Akt1^Y350F^ mice treated with Su/Hx exhibited an attenuated RV/LV+S ratio than treated WT mice (Fig.1C). The scatter plot demonstrates a strong positive correlation between RV/LV+S and RVSP across all groups (Fig.1D), indicating the consistency of these PAH indicators. It also shows that the Akt1^Y350F^ Su/Hx group is closer among untreated groups, indicating the resistance of Akt1^Y350F^ mice to PH.

H&E staining of lung sections, visually demonstrating differences in vascular remodeling, showed that WT mice treated with Su/Hx produce thickened vessel walls and narrowed lumens, while Akt1^Y350F^ mice treated with Su/Hx exhibit less severe vascular remodeling (Fig.1E). Occlusion score quantification of these observations (Fig.1F) indicate that Su/Hx treatment significantly increased the occlusion score in WT mice but Akt1^Y350F^ mice treated with Su/Hx showed a significantly lower occlusion score compared to treated WT mice (p < 0.05).

We elucidated the molecular mechanisms by which Akt1 nitration promotes pulmonary vascular remodeling, with a focus on endothelial-to-mesenchymal transition (EndMT) and proliferative signaling. Western blot analysis of TWIST1, a transcription factor known as a master regulator of EndMT ^14^, exhibits a significant increase in the Su/Hx group (Fig.2A). Akt1^Y350F^ mice showed a markedly attenuated TWIST1 response to Su/Hx treatment, with levels at the baseline and significantly lower than in Su/Hx WT mice. This suggests that Akt1 nitration is crucial for upregulating TWIST1 in response to PH-inducing conditions. Fig.2B shows the data on HIF1α expression, a key transcription factor involved in the cellular response to hypoxia that has been implicated in vascular remodeling in PAH ^15^. Similar to TWIST1, HIF1α levels were significantly elevated in Su/Hx-treated WT mice. Akt1^Y350F^ mice, however, showed a blunted HIF1α response to Su/Hx treatment, with levels significantly lower than in treated WT mice. This suggests that Akt1 nitration may contribute to the stabilization or upregulation of HIF1α under hypoxic conditions. HIF1α activates EndMT and the expression of alpha-smooth muscle actin (aSMA) in response to cellular stress and hypoxia ^16^. Our results showed that αSMA, a typical EndMT marker, was significantly upregulated in Su/HX-WT mice compared to the Su/Hx-Akt1^Y350F^ mice (Fig.2C), suggesting reduced EndMT in the mutant mice. STAT3, is involved in cell proliferation and survival, and its activation has been associated with EndMT and PH progression ^17,18^. In WT mice, Su/Hx treatment significantly increased p-STAT3 levels (Fig.2D). However, the total STAT3 levels have also increased, meaning it is regulated mainly by transcription rather than kinase signaling. Interestingly, Akt1^Y350F^ mice showed lower baseline p-STAT3 levels and a diminished response to Su/Hx treatment, with p-STAT3 levels significantly lower than in treated WT mice. This suggests that Akt1 nitration may enhance both p-STAT and STAT3 protein levels in PH.

**Figure 2.**
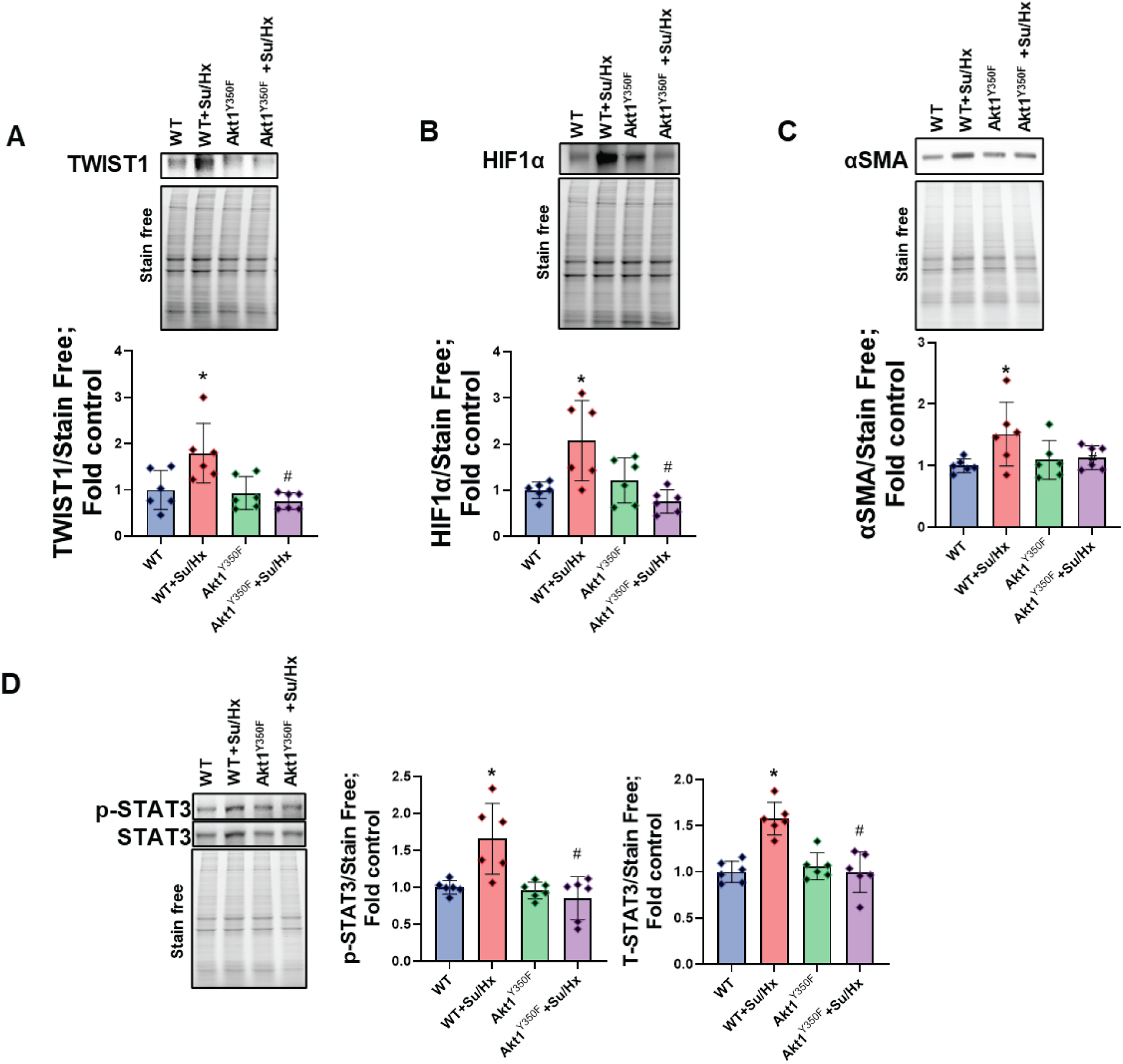
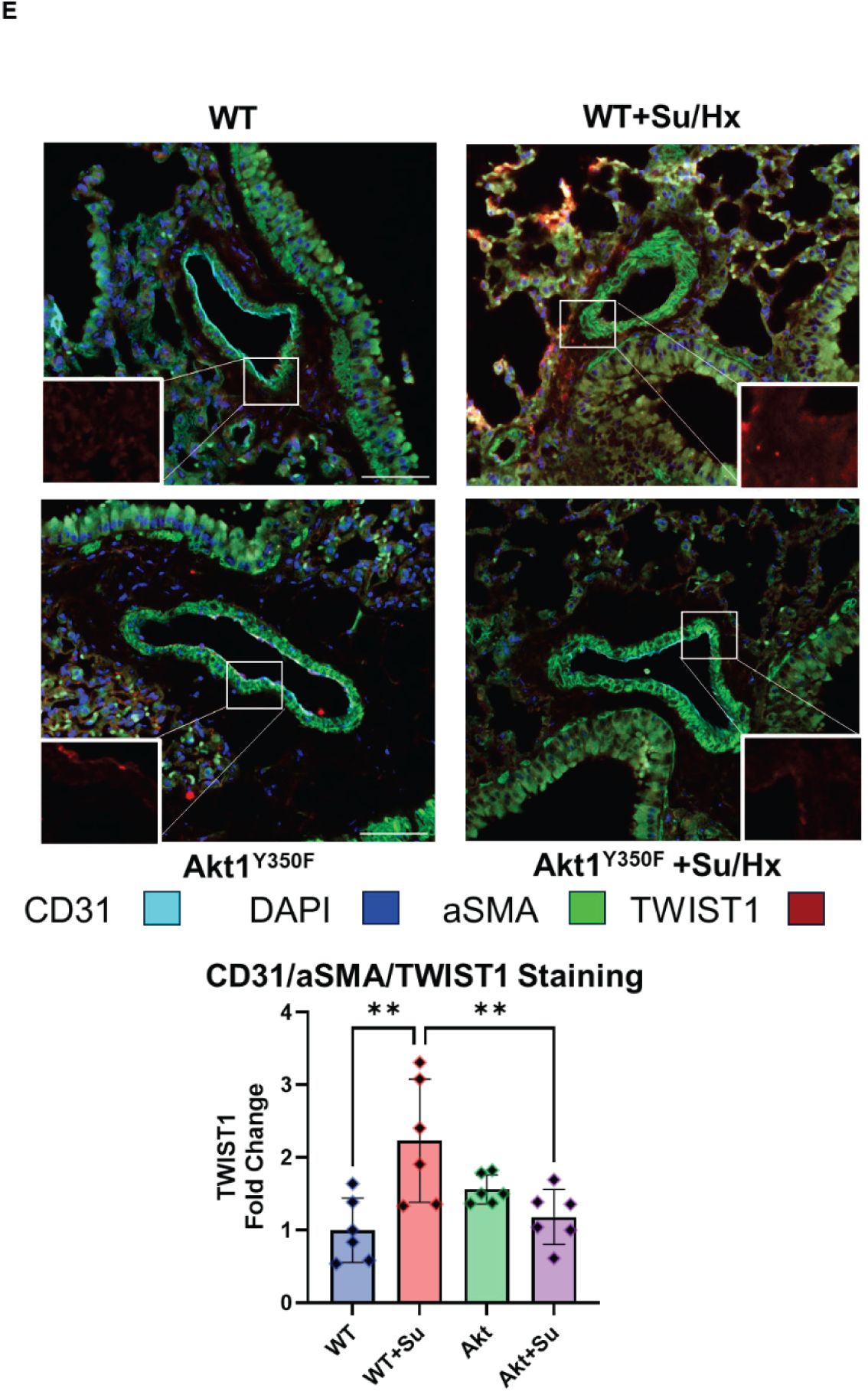
EndMT and Proliferative Markers: In the wild-type (WT) PAH model, the endothelial-to-mesenchymal transition marker TWIST1 (Figure 2A) and the hypoxia-inducible factor HIF1α (Figure 2B) were significantly increased. This increase was found to the activation of the proliferative marker STAT3 signaling (Figure 2C). However, the Akt1^Y350F^ model provided protection against EndMT and proliferative changes. Data are represented as Mean±SD, N=6, *p<0.05 vs. WT, ^#^p<0.05 vs. WT Su/HX by 1-way ANOVA with Bonferroni multiple comparison test.

Consistent with these observations, triple immunofluorescence staining for CD31, αSMA, and TWIST1 was performed in lung sections from Su/Hx WT and Akt1Y350F mice (Fig.2E). Quantitative analysis revealed that TWIST1 expression was significantly increased in Su/Hx WT lungs compared with Akt1Y350F mice, confirming the western blot data. Visually, CD31 staining appeared reduced in Su/Hx WT lungs relative to Akt1Y350F, suggesting a loss of endothelial marker expression under PH conditions. In contrast, αSMA staining showed comparable levels between the groups. These findings reinforce the conclusion that preventing Akt1 nitration attenuates TWIST1 induction and thereby limits EndMT and vascular remodeling in PH.

Using the lungs of WT and Akt1^Y350F^ mice, we performed a comprehensive proteomic analysis to investigate the effects of Akt1 nitration on protein dysregulation. This figure provides valuable insights into the broader cellular pathways affected by Akt1 nitration, expanding our understanding beyond the specific markers of EndMT examined in Figure 2. The proteomic data are organized into five categories, each representing the altered cellular functions relevant to PH pathogenesis. The data reveal significant changes in several proteins crucial for vascular development and remodeling (Fig.3A). Notably, endothelial nitric oxide synthase (eNOS) shows reduced expression in wild-type (WT) mice treated with SU5416/hypoxia (Su/Hx), while Akt1^Y350F^ mice maintain higher levels of eNOS. This finding is particularly relevant given the well-established role of nitric oxide in maintaining vascular homeostasis and its dysregulation in PAH ^19^. Other angiogenesis and remodeling-related proteins, such as fibronectin and chondroitin sulfate proteoglycan 4, show increased expression in WT Su/Hx-treated mice but are less elevated in Akt1^Y350F^ mice. These changes suggest that Akt1 nitration influences the angiogenic response in PAH, potentially contributing to the aberrant vascular remodeling observed in this condition. The proteomic analysis reveals significant changes in several enzymes involved in lipid synthesis, modification, and degradation (Fig.3B). For instance, 3-ketodihydrosphingosine reductase and lecithin retinol acyltransferase show increased expression in WT Su/Hx-treated mice, with a less pronounced increase in Akt1^Y350F^ mice. Conversely, some lipid metabolism enzymes, such as acetyl-coenzyme A synthetase 2-like, show decreased expression in the PAH model, attenuating this decrease in Akt1^Y350F^ mice. These findings suggest a potential connection between Akt1 nitration and altered lipid metabolism in PAH, which may impact signaling lipid production and energy metabolism in the diseased pulmonary vasculature. PH is known to be associated with mitochondrial dysfunction ^20^; Fig.3C provides insight into the mitochondrial metabolic changes related to PH and the influence of Akt1 nitration. Several mitochondrial proteins, including ferredoxin-2, carbonyl reductase 2 and coenzyme-A-related enzymes, show altered expression in the Su/Hx model, with reduced differences between WT and Akt/Su/Hx mice. These changes suggest that Akt1 nitration may impact mitochondrial function, potentially affecting energy production and oxidative stress in pulmonary vascular cells. The differences observed between WT and Akt1^Y350F^ mice indicate that preventing Akt1 nitration may help maintain mitochondrial homeostasis under PH conditions. Proteins involved in maintaining cell-cell communication are underexplored in PH pathogenesis and Fig.3D shows significant changes in their expressions. For example, integrin alpha-L and fascin actin-bundling protein 1 show decreased expression in WT Su/Hx-treated mice, but not in Akt1^Y350F^ mice. Proliferative c-Kit receptor exhibited high upregulation in the Su/Hx model, whereas Akt1/Su/Hx showed similar expression to

**Figure 3.**
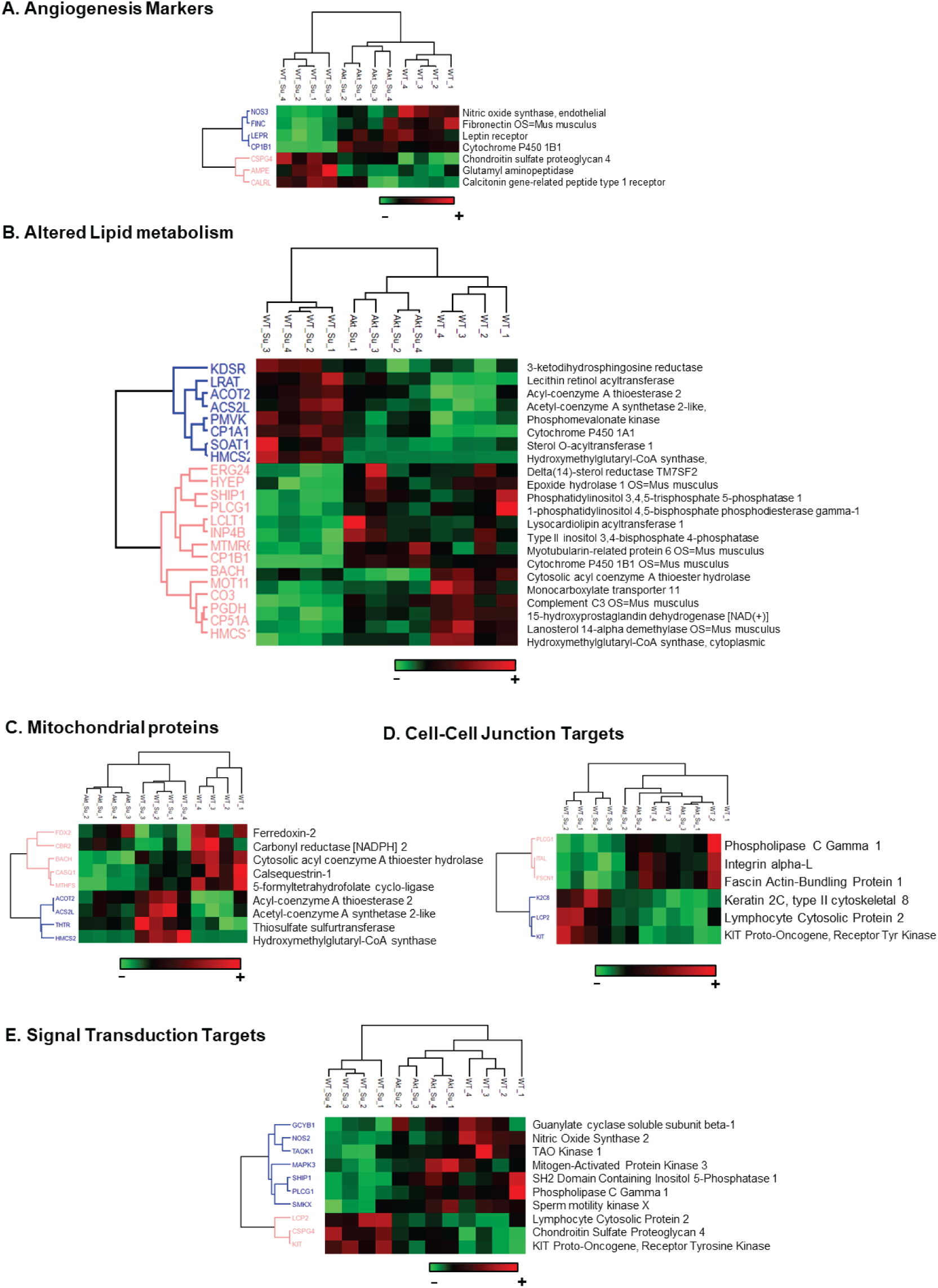
Proteomics Analysis: Lung proteomics analysis showed that Sugen hypoxia treatment in the wild-type (WT) model induced significant alterations in angiogenesis markers (Figure 3A), lipid metabolism (Figure 3B), mitochondrial function proteins (Figure 3C), cell-cell junction proteins (Figure 3D), and cellular signal transduction targets (Figure 3E). However, the Akt1^Y350F^ model helped to attenuate these proteomic changes. N=4. Gene ontology and Kyoto Encyclopaedia of Genes and Genomes (KEGG) pathway analysis were performed using DAVID, and the heat maps were generated using the Perseus bioinformatics platform. WT-Su represents WT sugen hypoxia and Akt-Su represents Akt1Y350F treated with sugen hypoxia.

WT. These changes suggest that Akt1 nitration may influence endothelial barrier integrity and intracellular communication, potentially contributing to the vascular leak and remodeling observed in PAH. Finally, Fig.3E focuses on signal transduction targets, providing a broader view of the signaling pathways affected by Akt1 nitration in PAH. The data show alterations in several key signaling molecules, including guanylate cyclase soluble subunit beta-1, mitogen-activated protein kinase 3, and phospholipase C gamma 1. The differential expression of these proteins between WT and Akt1^Y350F^ mice in the Su/Hx model suggests that Akt1 nitration has wide-ranging effects on cellular signaling cascades, potentially influencing processes such as vasodilation, cell proliferation, and calcium signaling.

To directly assess the role of Akt1 nitration in EndMT, we isolated primary ECs from WT and Akt1^Y350F^ mice and exposed them to TGF-β1 for 72 hours. WT ECs underwent a pronounced mesenchymal transition, acquiring an elongated, spindle-like morphology ^21^ (Fig. 4A). In striking contrast, Akt1^Y350F^ ECs maintained their endothelial phenotype under identical conditions, indicating resistance to TGF-β1-driven EndMT. At the molecular level, mesenchymal markers αSMA and SM22α ^22^ were significantly upregulated in WT ECs but remained unchanged in Akt1^Y350F^ cells (Fig. 4B**&C**). These findings establish Akt1 nitration as a critical determinant of EndMT induction in pulmonary vascular endothelium.

**Figure 4.**
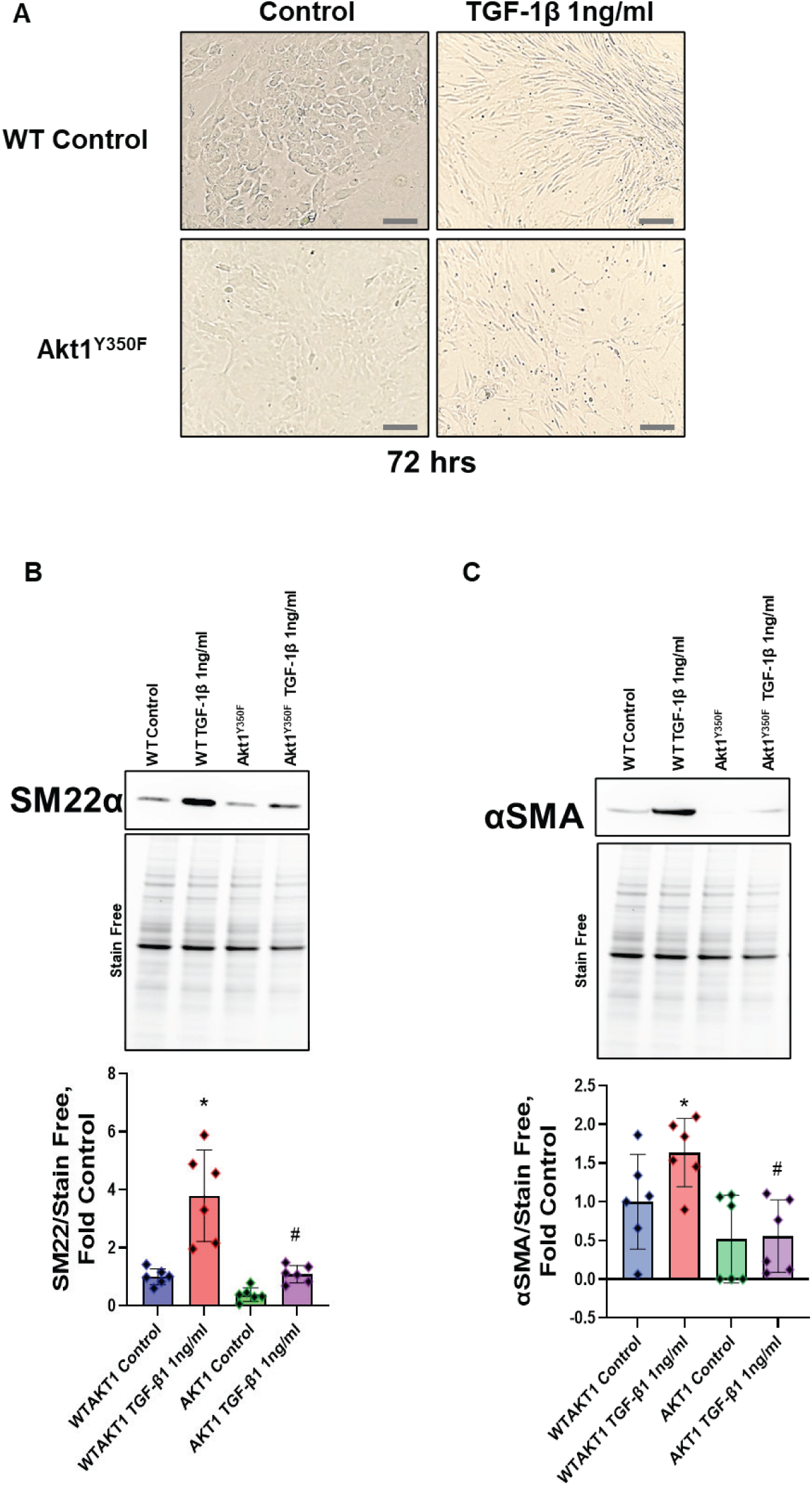
Akt1 nitration is required for TGF-β1induced EndMT in ECs. (Figure 4A) Representative phase-contrast images of ECs isolated from WT and Akt1^Y350F^ mice after 72 h of TGF-β1 stimulation. WT ECs adopt an elongated, spindle-shaped morphology indicative of EndMT, whereas Akt1^Y350F^ ECs retain cobblestone like endothelial morphology. (Figure 4B,C) Immunoblot analysis and quantification of mesenchymal markers αSMA (Figure 4B) and SM22α (Figure 4C) in WT and Akt1Y350F ECs following TGF-β1 treatment. Both proteins are significantly upregulated in WT ECs but attenuated in Akt1^Y350F^ ECs. Data are presented as Mean±SD, N=6, *p<0.05 vs. WT, ^#^p<0.05 vs. WT Su/HX by 1-way ANOVA with Bonferroni multiple comparison test.

## Discussion

This study provides convincing evidence for the critical role of Akt1 nitration in the pathogenesis of PAH. Using a novel Akt1^Y350F^ mutant mouse model resistant to nitration, we demonstrate that preventing Akt1 nitration significantly attenuates the development and progression of PH in the Su/Hx model. Our findings reveal multiple mechanisms through which Akt1 nitration contributes to the pathological vascular remodeling characteristic of PAH, offering new insights into the molecular foundation of this devastating disease. Our results show that Akt1 nitration significantly increases in the SU5416/hypoxia (Su/Hx) model of PH, and recent reports have also shown that Akt nitration is upregulated in PH patients ^23,24^. Preventing this nitration through the Y350F mutation substantially protects against PH development. This protection is evidenced by reduced right ventricular systolic pressure, decreased right ventricular hypertrophy, and diminished vascular occlusion in Akt1^Y350F^ mice subjected to Su/Hx treatment. These findings suggest that Akt1 nitration is not merely a consequence of PH but a critical driver of disease progression. The protective effects observed in Akt1^Y350F^ mice align with previous studies implicating Akt hyperactivation in PAH pathogenesis ^23^. However, our work extends beyond these earlier findings by identifying Akt1 nitration as a critical post-translational modification driving pathological events. This distinction is crucial, providing a more precise target for potential therapeutic interventions.

The molecular analyses reveal several key mechanisms through which Akt1 nitration promotes PH progression. The marked attenuation of TWIST1 upregulation in Akt1^Y350F^ mice under Su/Hx conditions suggests that Akt1 nitration is a critical mediator of EndMT in PAH pathobiology ^25^. This finding provides a mechanistic link between Akt1 nitration and the phenotypic changes in endothelial cells contributing to vascular remodeling (Fig.5). The blunted HIF1α response in Akt1^Y350F^ mice indicates that Akt1 nitration plays a role in amplifying the cellular response to hypoxia ^26^. This may explain, in part, the exaggerated vascular remodeling observed in PH under hypoxic conditions. Our observation of reduced STAT3 phosphorylation and total STAT3 levels in Akt1^Y350F^ mice suggests that Akt1 nitration enhances STAT3 signaling in PAH. Given STAT3’s known roles in promoting cell proliferation and survival ^18^, this finding provides insight into how Akt1 nitration may drive the hyperproliferative phenotype of pulmonary vascular cells in PAH. Importantly, all three transcriptional factors are also highly involved in EndMT. These molecular changes align well with the physiological and histological improvements observed in Akt1^Y350F^ mice, providing a mechanistic explanation for the protective effects of preventing Akt1 nitration.

**Figure 5.**
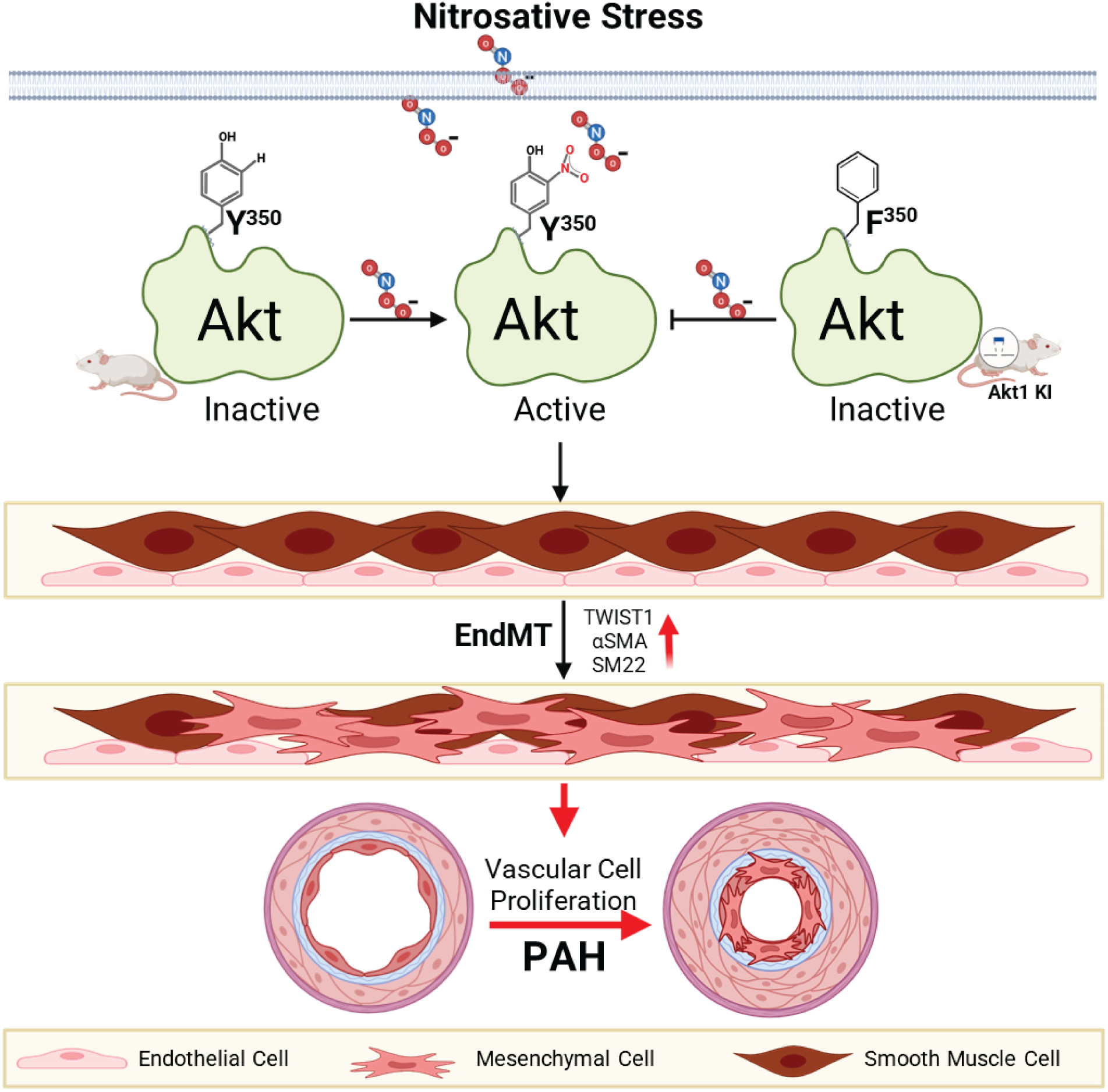
Akt1 Nitration as a Critical Driver of PAH Pathogenesis: Nitration of Akt1 at the tyrosine 350 residue can trigger Akt signaling activation, leading to endothelial-to-mesenchymal transition with TWIST1 and HIF1α. This results in altered cell signaling and vascular cell proliferation. Altogether, these changes contribute to pulmonary vascular remodeling and the development of PAH.

The proteomic analysis of lung tissue has uncovered additional pathways affected by Akt1 nitration in PAH, broadening our understanding of its impact. The differential expression of angiogenesis/remodeling-related proteins between WT and Akt1^Y350F^ mice suggests that Akt1 nitration influences angiogenic/remodeling response. This may contribute to the aberrant vascular proliferation characteristic of the disease. The alterations in lipid metabolism enzymes highlight a potential link between Akt1 nitration and metabolic reprogramming in PAH ^27^. This finding aligns with growing evidence of metabolic dysfunction in PAH pathogenesis ^28^. The changes observed in mitochondrial proteins suggest that Akt1 nitration may impact energy production and oxidative stress in pulmonary vascular cells. The receptors and junction protein alterations indicate that Akt1 nitration may affect endothelial barrier integrity and intercellular communication. This could contribute to the vascular leak and inflammation observed in PAH ^29^. Chondroitin Sulfate Proteoglycan 4 (CSPG4), also known as NG2, is a well-established marker for pericytes ^30^, critical microvasculature components. Our proteomic analysis revealed attenuation of this effector in Akt1^Y350F^ mice. This finding suggests a potential role for pericytes in the vascular remodeling process of PH, particularly in small arterioles. Pericytes are crucial in vascular stability, blood flow regulation, and angiogenesis. In the context of PH, the increased expression of CSPG4 could indicate enhanced pericyte proliferation or activation, which could contribute to the thickening of vessel walls, particularly in small arterioles, a hallmark of PH. It might also suggest altered pericyte-endothelial cell interactions, which could disrupt vascular homeostasis, potentially leading to increased vascular permeability and abnormal angiogenesis ^31^. Additionally, activated pericytes might produce excess extracellular matrix components, contributing to the increased vascular stiffness observed in PH. Under pathological conditions, pericytes have been shown to differentiate into myofibroblast-like cells, potentially contributing to vascular fibrosis in PH ^32^. The fact that Akt1^Y350F^ mice showed a less pronounced increase in CSPG4 suggests that Akt1 nitration may play a role in pericyte activation or proliferation in PH. This provides a new perspective on the cellular complexity of vascular remodeling in PH, extending beyond endothelial cells and smooth muscle cells to include pericytes. The proteomic analysis also revealed significant upregulation of c-KIT (CD117) in the Su/Hx model, with Akt1^Y350F^ mice showing expression levels similar to untreated WT mice. This finding is particularly intriguing given c-KIT’s known roles in stem cell biology and cellular transformation. c-KIT activation has been associated with increased cellular plasticity ^33^. In PH, this could contribute to the phenotypic switch of endothelial cells towards a more proliferative, mesenchymal-like state (EndMT). Increased c-KIT signaling might also promote the recruitment and activation of stem/progenitor cells to the pulmonary vasculature, potentially contributing to vascular remodeling ^34^. c-KIT signaling promotes cell survival and proliferation, which could contribute to the hyperproliferative phenotype of pulmonary vascular cells in PH. Furthermore, c-KIT interacts with various angiogenic pathways, and its upregulation could contribute to the dysregulated angiogenesis observed in PH ^35^. Akt1^Y350F^ mice maintained c-KIT expression levels similar to untreated controls, suggesting that Akt1 nitration may be a key mediator of c-KIT upregulation in PH. This provides a potential mechanistic link between Akt1 nitration and the transformation of endothelial cells into a more proliferative phenotype. The differential expression of CSPG4 and c-KIT in the examined model provides new insights into the cellular complexity of PH pathogenesis. Our findings highlight that PH involves endothelial and smooth muscle cells, pericytes, and potentially stem/progenitor cells. The fact that both CSPG4 and c-KIT expression were normalized in Akt1^Y350F^ mice suggests that Akt1 nitration may serve as a master regulator of multiple cellular processes in PH. Understanding the roles of CSPG4 and c-KIT in PH progression could lead to new therapeutic targets, potentially allowing for more precise interventions that address specific aspects of vascular remodeling. These proteomic findings support our primary observations and reveal the multifaceted impact of Akt1 nitration on pulmonary vascular cell biology in PAH.

Our findings support and extend previous studies on Akt signaling in PAH ^36^. While earlier work has implicated Akt hyperactivation in PAH progression ^37,38^, our study explicitly identifies Akt1 nitration as a critical post-translational modification driving this pathological activation. This nuanced understanding may help reconcile some conflicting reports on the role of Akt in PAH, as it suggests that the specific nature of Akt activation (i.e., nitration-induced) may be more important than overall Akt activity levels. The identification of Akt1 nitration as a key driver of PAH progression opens new avenues for therapeutic intervention. Strategies to prevent or reverse Akt1 nitration could stop or even reverse PAH progression. This could involve direct approaches to prevent Akt1 nitration via antioxidant peptide ^23^ or indirect methods to reduce overall nitrosative stress in the pulmonary vasculature. Moreover, our findings suggest that targeting the downstream effectors of nitrated Akt1, such as TWIST1, HIF1α, or STAT3, might offer alternative therapeutic strategies. The broad impacts of Akt1 nitration revealed by our proteomic analysis also suggest the potential for combination therapies simultaneously addressing multiple aspects of vascular proliferation.

While our study provides significant insights, several limitations and areas for future research should be acknowledged. Our findings are based on a mouse Su/Hx model, which is not a severe model of PH ^39^. Further validation will require translational approaches. Additional research is needed to elucidate the precise mechanisms leading to increased Akt1 nitration in PAH. Our study focused on whole-lung analyses. Future work should investigate the cell-specific effects of Akt1 nitration in different pulmonary vascular cell types. Long-term studies are needed to assess the durability of the protective effects conferred by preventing Akt1 nitration via antioxidant peptides with affinity to Akt. While our findings suggest therapeutic potential, significant work remains to translate interventions targeting Akt1 nitration or its downstream effects to clinics.

In conclusion, our study identifies Akt1 nitration as a critical driver of PAH pathogenesis, influencing multiple aspects of pulmonary vascular cell biology. By demonstrating the protective effects of preventing Akt1 nitration, we provide new insights into the molecular mechanisms underlying PAH and suggest novel therapeutic strategies for this devastating disease (Fig.5). As we continue to unravel the complex pathways involved in PAH, targeting Akt1 nitration may offer a promising approach to halt or reverse disease progression, potentially improving outcomes for patients with this challenging condition.

## Methods

### Animal Models

#### Generation of Akt1^Y350F^ mice

Akt1^Y350F^ knock-in (KI) mice on a C57BALB/c background were generated using CRISPR/Cas9 technology by the GEMM (Genetically Engineered Mouse Models) Core at the UA BIO5 Institute, University of Arizona (Tucson, AZ). This mouse model was created by introducing a single-point mutation, substituting tyrosine with phenylalanine at residue 350. Genotyping of the offspring was performed using PCR with the following primers: forward primer (CACATCAAGATAACGGACTTCG) labeled with FAM (6-fluorescein amidite) and reverse primer (CTGTGTAGGGTCCTTCTTGAGC), provided by the University of Arizona Genetics Core.

Previously, we reported that nitration of the Akt protein at the tyrosine 350 site can activate Akt signaling, contributing to the pathogenesis of pulmonary arterial hypertension (PAH) ^23^. To further explore the mechanistic effects of Akt nitration, we used female mice aged 10 ± 1 weeks (n = 6–9). The mice were housed at 22°C on a 12-hour light/dark cycle with access to standard rodent chow and water. All experimental protocols were approved by the Indiana University Institutional Animal Care and Use Committee. To induce pulmonary arterial hypertension (PAH), animals were administered with Sugen-5416 (20 mg/kg/week, intraperitoneally (IP)) in combination with hypoxia (10% O_2_) for four weeks. The study included four groups: control, Sugen/Hypoxia (Su/Hx), Akt control, and Akt with Su/Hx for 4 weeks.

### Hemodynamic Measurement

Mice were anesthetized with a combination of Ketamine (100 mg/kg) and Xylazine (10 mg/kg) from MWI (501072 and 510004, Boise, ID), administered IP. Right ventricular systolic pressure (RVSP) was recorded using an SPR-1000 catheter (Millar Instruments, Houston, TX). To measure RVSP, the catheter was inserted via the right jugular vein and advanced into the right ventricle (RV). Hemodynamic data were collected using the AD Instruments PowerLab-4/35 system (Colorado Springs, CO).

Following these measurements, the mice were connected to a MiniVent ventilator Type-845 from Harvard Apparatus (South Natick, MA) via a tracheal catheter. The lungs were perfused with a 0.9% NaCl solution through the RV. Afterward, the lungs and heart were excised, and the Fulton index was calculated by measuring the ratio of the RV free wall to the combined weights of the left ventricle (LV) and septum (S). The left lung was fixed in formalin for histological examination, and the remaining lung tissue was stored at −80°C for protein expression studies.

### Histopathological Analysis

For the morphometric evaluation, 5 μm cut tissue sections were deparaffinized and stained with H&E (hematoxylin and eosin) following the standard procedures of the TACMASR (Tissue Acquisition and Cellular/Molecular Analysis Shared Resource) at the Cancer Center, University of Arizona (Tucson, AZ). Six to seven randomly selected pulmonary arteries (PAs) per animal, with diameters ranging from 50 μm to less than 300 μm, were examined using a 40X objective lens on an Echo RVL2-k2 Revolve Fluorescence Microscope (San Diego, CA). The morphometric analysis was conducted in a blinded manner to ensure unbiased results. All histological and microscopy procedures were carried out concurrently under identical conditions. Pulmonary artery occlusion score was quantified using Fiji ImageJ software (Version 1.52p, National Institutes of Health, Washington; http://fiji.sc/Fiji) ^40^.

### Protein Analysis

To analyze total protein content, lung tissue lysate was prepared by homogenizing in a tissue permeabilization buffer containing protease and phosphatase inhibitors (Halt™ Protease and Phosphatase Inhibitor Cocktail) from Thermo Fisher Scientific (Rockford, IL). The tissue or cell homogenates was centrifuged at 10,000 g for 10 minutes, and the supernatant was used for protein quantification using the Pierce BCA Protein Assay Kit (Thermo Fisher Scientific, Burlington, ONT). Samples were prepared with 6X Laemmli buffer (Boston Bioproducts Inc., Ashland, MA) and heated at 95°C for 5 minutes. Gel electrophoresis was performed using a 4–20% Stain-Free gel (Bio-Rad Laboratories Inc., Hercules, CA). The proteins were then transferred to a polyvinylidene difluoride (PVDF) membrane with Trans-Blot Turbo System (Bio-Rad Laboratories Inc., Hercules, CA). Subsequently, the membranes were then blocked in EveryBlot Blocking Buffer (Bio-Rad Laboratories Inc., Hercules, CA) and incubated overnight with the following primary antibodies: custom-made Akt Y350 NO2 antibody (1:1,000, Pacific Immunology Corp., Ramona, CA, USA) (Varghese et al., 2020), TWIST1 (1:1,000, ab50581, Abcam, Waltham, MA), Phospho-Stat 3 (Ser^727^) (1:1,000, 9139, CST), and Stat 3 (1:1,000, 9139, CST, Danvers, MA).

The reactive bands were imaged with the Bio-Rad ChemiDoc-MP System, and band intensity was analyzed using Image Lab software. Protein expression was assessed with stain-free gel images and are then normalized to total protein levels, following the method described by Rivero-Gutierrez et al. (2014) ^41^. Certain membranes were reused by stripping with Restore Western Blot Buffer (21063, Thermo Fisher Scientific, Rockford, IL) and reprobed to detect additional target proteins.

### Proteomic Analysis

#### In-gel digestion

For the proteome wide experiments, 50 mg of mouse whole lung lysate was separated by SDS-PAGE, and each lane was cut into five slices. The gel slices were subjected to trypsin digestion and the resulting peptides were purified by C^18^-based desalting exactly as previously described ^42,43^. In brief, the SDS-PAGE gel slices were placed in a 0.6 mL LoBind polypropylene tube (Eppendorf), destained twice with 375 mL of 50% acetonitrile (ACN) in 40 mM NH_4_HCO_3_ and dehydrated with 100% acetonitrile (CAN) for 15 minutes. After removal of the ACN by aspiration, the gel pieces were dried in a vacuum centrifuge at 60°C for 30 minutes. Trypsin (250 ng; Sigma-Aldrich) in 20 mL of 40 mM NH_4_HCO_3_ was added, and the samples were maintained at 4°C for 15 minutes prior to the addition of 50-100 mL of 40 mM NH_4_HCO_3_. The digestion was allowed to proceed at 37°C overnight and was terminated by addition of 10 mL of 5% formic acid (FA). After further incubation at 37°C for 30 minutes and centrifugation for 1 minute, each supernatant was transferred to a clean LoBind polypropylene tube. The extraction procedure was repeated using 40 mL of 0.5% FA, and the two extracts were combined and dried down to ∼5-10 mL followed by the addition of 10 mL of 0.05% heptafluorobutyric acid/5% FA (vol/vol) and incubation at room temperature for 15 minutes. The resulting peptide mixtures were loaded on a solid phase C18 ZipTip (Millipore, Billerica, MA) and washed with 35 mL 0.005% heptafluorobutyric acid/5% FA (vol/vol) followed by elution first with 4 mL of 50% ACN/1% FA (vol/vol) and then a more stringent elution with 4 mL of 80% ACN/1% FA (vol/vol). The eluates were combined and dried completely by vacuum centrifugation and 6 mL of 0.1% FA (vol/vol) was added followed by sonication for 2 minutes. 2.5 mL of the final sample was then analyzed by mass spectrometry.

#### Mass spectrometry and database search

HPLC-ESI-MS/MS was performed in positive ion mode on a Thermo Scientific Orbitrap Fusion Lumos tribrid mass spectrometer fitted with an EASY-Spray Source (Thermo Scientific, San Jose, CA) as previously described ^43^. In brief, NanoLC was performed using a Thermo Scientific UltiMate 3000 RSLCnano System with an EASY Spray C18 LC column (Thermo Scientific, 50 cm x 75 mm inner diameter, packed with PepMap RSLC C18 material, 2 mm, cat. # ES803); loading phase for 15 minutes; mobile phase, linear gradient of 1–47% ACN in 0.1% FA for 106 minutes, followed by a step to 95% ACN in 0.1% FA over 5 minutes, hold 10 minutes, and then a step to 1% ACN in 0.1% FA over 1 minute and a final hold for 19 minutes (total run 156 minutes); Buffer A = 100% H_2_O in 0.1% FA; Buffer B = 100% ACN in 0.1% FA; flow rate, 300 nL/min. All solvents were liquid chromatography mass spectrometry grade. Spectra were acquired using XCalibur, version 2.1.0 (Thermo Scientific). A “top 15” data-dependent MS/MS analysis was performed (acquisition of a full scan spectrum followed by collision-induced dissociation mass spectra of the 15 most abundant ions in the survey scan). Dynamic exclusion was enabled with a repeat count of 1, a repeat duration of 30 seconds, an exclusion list size of 500, and an exclusion duration of 40 seconds Tandem mass spectra were extracted from Xcalibur ‘RAW’ files and charge states were assigned using the ProteoWizard 2.1.x msConvert script using the default parameters. The fragment mass spectra were searched against the 2021 *Homo sapiens* SwissProt database (20376 entries) using Mascot (Matrix Science, London, UK; version 2.4) using the default probability cut-off score. The search variables that were used were: 10 ppm mass tolerance for precursor ion masses and 0.5 Da for product ion masses; digestion with trypsin; a maximum of two missed tryptic cleavages; variable modifications of oxidation of methionine and phosphorylation of serine, threonine, and tyrosine. Cross-correlation of Mascot search results with X! Tandem was accomplished with Scaffold (version Scaffold 4.8.7; Proteome Software). Probability assessment of peptide assignments and protein identifications were made using Scaffold. Only peptides with ≥ 95% probability were considered.

#### Label-free Quantitative Proteomics

Progenesis QI for proteomics software (version 2.4, Nonlinear Dynamics Ltd., Newcastle upon Tyne, UK) was used to perform ion-intensity based label-free quantification similar to as previously described (PMID 31018989). In brief, in an automated format, .raw files were imported and converted into two-dimensional maps (y-axis = time, x-axis =m/z) followed by selection of a reference run for alignment purposes. An aggregate data set containing all peak information from all samples was created from the aligned runs, which was then further narrowed down by selecting only +2, +3, and +4 charged ions for further analysis. A peak list of fragment ion spectra was exported in Mascot generic file (.mgf) format and searched against the 2022 *Mus musculus* SwissProt database (17138 entries) using Mascot (Matrix Science, London, UK; version 2.6). The search variables that were used were: 10 ppm mass tolerance for precursor ion masses and 0.5 Da for product ion masses; digestion with trypsin; a maximum of two missed tryptic cleavages; variable modifications of oxidation of methionine and phosphorylation of serine, threonine, and tyrosine; 13C=1. The resulting Mascot .xml file was then imported into Progenesis, allowing for peptide/protein assignment, while peptides with a Mascot Ion Score of <25 were not considered for further analysis. Precursor ion-abundance values for peptide ions were normalized to all proteins. Unbiased hierarchal clustering analysis (heat map) was performed in Perseus ^44,45^. Gene ontology and KEGG pathway enrichment analysis was performed with DAVID ^46^.

### Immunofluorescence Staining

Formalin-fixed, paraffin-embedded (FFPE) mouse lung tissues were sectioned at a thickness of 4 µm using a rotary microtome and mounted onto charged glass slides. Tissue sections were deparaffinized in xylene and rehydrated through a graded ethanol series, followed by heat-induced epitope retrieval in citrate buffer (pH 6.0) using a pressure cooker for 10 minutes. After cooling to room temperature, slides were washed in phosphate-buffered saline (PBS) and blocked in 5% donkey serum containing 0.1% Triton X-100 for 1 hour to reduce nonspecific binding.

Triple immunofluorescence staining was performed at the Histology Core Facility, Indiana University School of Medicine, using primary antibodies against CD31 (goat anti-mouse), αSMA (mouse anti-mouse), and TWIST1 (rabbit anti-mouse). Sections were incubated overnight at 4°C with primary antibodies diluted in blocking buffer. After PBS washes, species-specific secondary antibodies conjugated to fluorophores were applied for 1 hour at room temperature: donkey anti-goat Alexa Fluor 555, donkey anti-mouse Alexa Fluor 488, and donkey anti-rabbit Alexa Fluor 647. Nuclei were counterstained with DAPI (1 µg/mL) for 5 minutes. Slides were washed, mounted in antifade mounting medium, and cover slipped.

Fluorescence imaging was performed using an Echo Revolve fluorescence microscope with consistent acquisition settings across all samples. For each biological replicate (n = 6), 6 to 8 non-overlapping fields were imaged per lung section, focusing on representative areas of the parenchyma. Image analysis and quantification were conducted using FIJI (ImageJ, NIH). Quantitative parameters, including mean fluorescence intensity, area fraction, and co-localization metrics, were measured. All image processing and analyses were performed in a blinded fashion to ensure unbiased interpretation.

### Isolation of Endothelial Cells

Whole lungs from wild-type (WT; C57BL/6) and Akt1^Y350F^ KI mice were used for endothelial cell (EC) isolation via magnetic-activated cell sorting, following previously described protocol^47^. Briefly, 12 to 13-week-old WT and Akt1^Y350F^ mice (n = 5 per group) were euthanized under anesthesia. Pulmonary perfusion was performed using 0.9% saline via the RV to eliminate blood contaminants. The lungs were then excised, minced, and enzymatically digested with a cocktail containing 1 mg/mL neutral protease (LS02109, Worthington, Lakewood, NJ) and collagenase type II (LS004176, Worthington, Lakewood, NJ) for 45 minutes at 37°C.

The resulting cell suspension was sequentially filtered through 70 µm (CLS431751, Corning, New York, USA) and 40 µm (CLS431750, Corning, New York, USA) cell strainers to obtain a single-cell suspension. Endothelial cells were isolated by incubating the suspension with anti-mouse CD31 microbeads (130-097-418, Miltenyi Biotec, Westphalia, Germany) to enrich for pulmonary artery endothelial cells (MPAECs) via magnetic separation. The sorted MPAECs were seeded onto 6-well plates and cultured in complete EC medium (1001, ScienCell, Carlsbad, CA) supplemented with 5% fetal bovine serum (FBS) and antibiotic-antimycotic solution (15240062, Thermo Fisher Scientific). Cells exhibited characteristic cobblestone morphology and were expanded up to passage 3 for experimental use.

### In vitro Endothelial to Mesenchymal Transition

To induce EndMT, confluent ECs from both experimental groups were treated with 1 ng/mL transforming growth factor-beta 1 (TGF-β1; PeproTech Inc., Cranbury, NJ, USA) for three days^48^, with media refreshed every other day. Control cells received culture medium without any cytokines. Morphological changes were monitored using Echo Revolve fluorescence microscope, and EndMT was confirmed by assessing protein expression of mesenchymal markers SM22α (40471, Cell Signaling Technology, Danvers, Massachusetts, USA) and αSMA (14-9760-82, Invitrogen, Waltham, MA, USA) via western blot analysis as described above.

### Statistical Analysis

Statistical analyses were performed with GraphPad Prism software - 10.3.1 version. For each sample, mean values with standard deviation (± SD) were calculated. Outliers were detected using Grubbs’ test through the GraphPad outlier tool (alpha=0.05). Significance was assessed using ANOVA (analysis of variance). For one-way ANOVA, Bonferroni’s post hoc test was used to compare selected pairs of columns. A 95% confidence interval was used to determine statistical significance.

## Abbreviations

ACN: Acetonitrile
Akt1^Y350F^: Akt1 with tyrosine 350 replaced by phenylalanine
ANOVA: Analysis of Variance
BCA: Bicinchoninic Acid
c-KIT: Tyrosine-protein kinase Kit (CD117)
CRISPR: Clustered Regularly Interspaced Short Palindromic Repeats
CSPG4: Chondroitin Sulfate Proteoglycan 4
DAVID: Database for Annotation, Visualization and Integrated Discovery
EndMT: Endothelial-to-Mesenchymal Transition
eNOS: Endothelial Nitric Oxide Synthase
FA: Formic Acid
FAM: 6-Fluorescein Amidite
H&E: Hematoxylin and Eosin
HIF1α: Hypoxia-Inducible Factor 1-alpha
HPLC-ESI-MS/MS: High-Performance Liquid Chromatography-Electrospray Ionization-Tandem Mass Spectrometry
IP: Intraperitoneally
KEGG: Kyoto Encyclopedia of Genes and Genomes
KI: Knock-in
LV: Left Ventricle
N-Akt: Nitrated Akt
NG2: Neural/Glial Antigen 2 (another name for CSPG4)
ONOO-: Peroxynitrite
PAH: Pulmonary Arterial Hypertension
PAs: Pulmonary Arteries
PCR: Polymerase Chain Reaction
PH: Pulmonary Hypertension
PVDF: Polyvinylidene Difluoride
RV: Right Ventricle
RVSP: Right Ventricular Systolic Pressure
RV/LV+S: Right Ventricle to Left Ventricle plus Septum ratio (Fulton index)
SDS-PAGE: Sodium Dodecyl Sulfate-Polyacrylamide Gel Electrophoresis
STAT3: Signal Transducer and Activator of Transcription 3
Su/Hx: Sugen-5416/Hypoxia
TWIST1: Twist-related protein 1
WT: Wild-type

## Acknowledgments

This work was supported by the following grants: National Institutes of Health (NIH) R01 grants HL133085 and HL160666 to O. Rafikova; NIH R01 grants HL132918 and HL151447 to R. Rafikov; NIH K99 grant 1K99HL171869 to J. James; the American Heart Association grant 969574 to O. Rafikova, 834220 to J. James, and 831538 to M.V. Varghese. The figures were created using elements of bio-render and Servier Medical Art.

## Authorship Contributions

Conception and design: RR, OR; Analysis and interpretation: JJ, MVV, DB, PL; Drafting the manuscript for important intellectual content: MVV, RR, OR

